# Constructed Negatives Shape the Evaluation of Sequence-Based Protein–Protein Interaction Models

**DOI:** 10.64898/2026.10.08.757593

**Authors:** Enzo Durand, Sergei Grudinin, Béatrice Durand, Élodie Laine

## Abstract

Many machine-learning tasks, including representation learning, rely on negative examples that are constructed rather than directly observed. This issue is especially important for protein–protein interaction (PPI) prediction, where experimentally established non-interactions are scarce. PPI models are therefore commonly trained and evaluated against sampled, unobserved protein pairs treated as negatives. We study how this construction affects model comparison. Keeping positives, protein-disjoint splits, class balance, and model selection fixed, we vary training and evaluation negatives independently across six constructions and four sequence-based models, three of which use protein language model (pLM) representations. Changing only the test negatives shifts AUPRC by up to 0.26, and the top-ranked architecture changes with the construction. Negatives matched to true partners in pLM representation space reduce all pLM-based models to near-chance performance, and shuffling partners within each class shows that the original pairing contributes little to AUROC. Constructed training negatives rarely outperform uniform sampling: on external data, they underperform it in 108 of 120 comparisons for the pLM-based models, with gains never exceeding 0.05 AUROC and losses up to 0.23. We release splits, negative-generation procedures, and evaluation code that allow training and evaluation negatives to be varied separately. Our results show that with constructed negatives, model comparisons are not just noisy but systematically sensitive to how negatives are defined, a methodological choice that is often left implicit.

## 1 Introduction

In many machine-learning tasks, the negative class cannot be observed directly and must instead be constructed, because most data sources record what occurred rather than what did not. Protein–protein interaction (PPI) prediction is a prominent instance of this problem. It is usually formulated as binary classification, where a model receives two protein sequences, typically encoded with representations from protein language models (pLMs), and predicts whether they interact. Positive examples can be collected from interaction databases and systematic assays, but experimentally established non-interactions are much rarer. Large labelled PPI datasets therefore usually construct the negative class from protein pairs that have not been reported as interacting.

This construction is necessary for large-scale supervised learning, but it changes the statistical problem given to the model. A negative generator can alter per-protein properties (degree, sequence length, amino-acid composition, taxonomic origin, frequency in the dataset), pair-level properties such as functional relatedness, and which regions of the pLM representation space are populated. From a machine-learning perspective, changing the generator changes the class-conditional input distribution and can create distribution shift between benchmark constructions. If these properties differ between positive and negative classes, a classifier can use them to distinguish the classes even when they provide little information about compatibility of the specific proteins in a pair, a form of shortcut learning. High classification performance may therefore reflect regularities introduced by dataset construction rather than pair-specific interaction information.

This problem has been studied from several directions. Negative-set design can make PPI classification artificially easy [Ben-Hur and Noble, 2006], and negative sampling for model training and performance estimation have different requirements [Park and Marcotte, 2011]. Protein reuse and sequence similarity across train and test sets can invalidate pair-input generalization claims [Park and Marcotte, 2012, Bernett et al., 2024]. More recent work has identified degree, taxonomy, representation-level similarity, and other dataset-construction shortcuts [Li et al., 2025, Hallee et al., 2026, Szymborski and Emad, 2026, Bernett et al., 2026]. Together, these studies motivate evaluating the negative construction used during training separately from the construction used during testing.

We address this problem with **PPIShift**, a controlled evaluation of sequence-based PPI models on the human Gold benchmark introduced by Bernett et al. [2024]. We hold fixed its positive interactions and protein-disjoint split, class balance, common 512-residue sequence subset, model-selection procedure, and random seeds. We evaluate four complementary sequence-based models: a controlled frozen-pLM pair multi-layer perceptron (MLP) baseline based on ESM-2 Lin et al. [2023], TUnA [Ko et al., 2024], RAPPPID [Szymborski and Emad, 2022], and D-SCRIPT-ESM2 [Reim et al., 2025]. For each model, the six training-negative constructions are crossed with the same six test-negative constructions, giving a 6 *×* 6 matrix per model and seed. This separates the effect of training negatives from the effect of test negatives. The six constructions range from uniform random pairing to degree matching, anchor-based matching of the replaced partner in sequence-feature or ESM-2 space, a global bias-reducing ILP [Bernett et al., 2026], and topology-driven sampling [Chatterjee et al., 2025].

We perform *cross-source* evaluations, applying the same Gold-trained models, without retraining or threshold adaptation, on independent data sources. Protein isoform-specific yeast two-hybrid (Y2H-iso) assay outcomes [Lambourne et al., 2026] and explicit curated IntAct negatives [Orchard et al., 2014] are kept distinct from BioGRID [Oughtred et al., 2021] and STRING [Szklarczyk et al., 2025] evaluations that pair source-specific positives with constructed negatives. Simple property-based controls, protein-only controls, and within-class partner shuffling test whether discrimination requires the specific protein pair. PPIShift therefore evaluates whether conclusions about a model remain consistent across negative constructions, partner-shuffling controls, and evaluation sources. Our goal is not to identify a universally preferable negative sampler, but to determine which conclusions about sequence-based PPI models, including comparisons between pLM-based and raw-sequence architectures, remain consistent when the negative class and evaluation source change.

Our main findings are as follows. (i) Performance and model rankings depend on both training and test negative construction. Anchor-based negatives matched in ESM-2 space are the hardest, with AUPRC close to chance for the ESM2-based models regardless of training construction, followed by sequence-matched and bias-reduced negatives; topology-driven negatives are the easiest. (ii) No construction removes all simple predictive signals, and partner shuffling shows that the original pairing contributes little to AUROC. (iii) Constructed training negatives rarely outperform uniform training outside their own construction. Training on anchor-matched negatives brings little benefit even on test sets built the same way, and models trained on negatives matched in ESM-2 space perform close to chance on every test set, including the easiest. Bias-reducing and topology-driven training improve transfer to some external sources for some models, but not consistently. (iv) The only model trained from raw sequence, RAPPPID, remains close to chance, consistent with a benefit of pretrained representations, although this comparison is confounded with architecture.

## 2 Related Work

Reliable evaluation is a broader problem across biological machine learning, where labels are expensive and train–test similarity can be strong. PINDER and PLINDER use similarity-aware splitting for protein-complex and protein–ligand evaluation [Kovtun et al., 2024, Durairaj et al., 2024]; PoseBusters shows that favorable docking metrics need not imply physically valid or sequence-novel predictions [Buttenschoen et al., 2024]; and Bohl et al. [2026] report that several domain-adaptation methods do not outperform simple baselines for single-cell drug-response transfer. These settings differ from binary PPI prediction but motivate the same separation between benchmark performance and robust generalization under controlled shifts.

### Sequence-based PPI models

Recent sequence-based PPI predictors use either trainable sequence encoders or representations from pretrained pLMs. TUnA uses ESM-2 residue representations with intra-protein and inter-protein Transformer encoders and an SNGP output layer [Ko et al., 2024, Lin et al., 2023]; RAPPPID uses SentencePiece tokenization and twin bidirectional AWD-LSTM encoders [Szymborski and Emad, 2022]; and D-SCRIPT-ESM2 denotes the ESM-2-based reimplementation evaluated by Reim et al. [2025], derived from the original D-SCRIPT architecture [Sledzieski et al., 2021]. We also include a controlled Frozen-ESM2 pair MLP baseline that symmetrically combines fixed ESM-2 (150M) single-protein representations. These models span pooled-pLM, residue-level interaction, Transformer/SNGP, and raw-sequence recurrent architectures while remaining feasible to retrain across the full factorial. Contemporary context includes C3PI, a sequence-based classifier evaluated on Gold [Hosseini et al., 2025], large-scale coevolution/structure-based human interactome prediction [Zhang et al., 2025], and the graph-level PRING benchmark at NeurIPS 2025 [Zheng et al., 2025]. The four models were fixed before final-test inspection, so no architecture was added post hoc.

### Dataset shortcuts

Distribution shift and leakage remain central issues in protein ML. FLIP2 explicitly evaluates protein models under application-relevant distribution shifts [Didi et al., 2026]; PRING introduces redundancy-and leakage-controlled PPI network evaluation [Zheng et al., 2025]; and Bernett et al. [2024] showed that protein sharing, sequence similarity, and interacting hubs can strongly affect sequence-based PPI evaluation. DataSAIL provides a general similarity-aware splitting framework [Joeres et al., 2025, 2026]. Pretrained representations create a separate exposure issue because downstream benchmark proteins or close relatives may occur in the pretraining corpus [Szymborski and Emad, 2026]. Taxonomic identity can also be recoverable from protein representations [Hallee et al., 2026]; our main Gold experiment is human-only, so we do not interpret taxonomy as a Gold-specific shortcut.

### Negative construction

Negative-example construction is itself an active methodological problem [Li et al., 2025]. Random pairing is scalable but can create positive-versus-negative differences that are predictive without encoding specific molecular compatibility. This issue was identified early for localization-informed negatives [Ben-Hur and Noble, 2006], and Park and Marcotte [2011] explicitly distinguished negative sampling for training from negative sampling for performance estimation. More recent approaches include topology-driven TPPNI [Chatterjee et al., 2025], bias-reducing ILP construction [Bernett et al., 2026], and reconstruction of experimentally tested search spaces to recover negative evidence [Ås et al., 2026]. Our contribution is complementary: we vary training and test negative constructions independently and ask whether model comparison, pair dependence, and cross-source performance change with that choice.

Our contributions are fourfold:

- a controlled training-negative *×* test-negative evaluation of four sequence-based PPI models across six constructions
- direct controls for pair-specific learning, including property-based baselines, protein-only prediction, and partner-shuffled evaluation
- evaluation of the same Gold-trained models on Y2H-iso assay outcomes and an IntAct set with curated negatives, cross-source BioGRID/STRING data, and a separate leakage-reduced DataSAIL split
- a reusable evaluation pipeline that allows new PPI architectures to be tested under the same constructions, controls, and evaluation rules.

## 3 Data, Models, and Negative-Pair Construction

### 3.1 Gold benchmark and external evaluation sources

Our controlled experiment starts from the human Gold benchmark introduced by Bernett et al. [2024] and subsequently used by Reim et al. [2025]. In the published Gold split, proteins are assigned to disjoint train (Intra1), validation (Intra0), and test (Intra2) partitions, so a protein appearing in one partition does not reappear in another. The source split contains 81,596 training, 29,630 validation, and 26,024 test positive pairs, with degree-balanced constructed negatives in the original release [Bernett et al., 2024, Joeres et al., 2026].

To evaluate all four models on identical raw sequences, we apply a predefined maximum of 512 amino-acid residues per protein before constructing candidate sets or negatives. This is the minimum native input limit of the fixed model implementations, not a threshold selected from test performance or desired sample size. The common subset retains 13,304 training, 11,085 validation, and 10,111 test positive pairs, corresponding to 16.3% of the original Gold training positives. All main Gold claims therefore apply to this sequence-compatible subset.

For each Gold split, the candidate set contains proteins appearing in the retained positive pairs. For an anchor protein and observed positive partner, a replacement candidate excludes the anchor, the observed partner, and any pair present in the fixed positive-pair exclusion registry for that stage. We generate one constructed negative per positive. Positive pairs are identical across constructions; only negative pairs change. A constructed Gold negative is therefore an *unobserved pair* under these exclusion rules, not an experimentally established non-interaction.

For external evaluation, we use a fixed sequence-compatible Y2H-iso subset with 846 detected interactions and 3,185 assay non-detections [Lambourne et al., 2026], and an IntAct set with 10,111 positive records and 236 explicit curated negatives [Orchard et al., 2014]. Y2H-iso non-detections are assay outcomes and may include interactions missed because of assay sensitivity or other technical limitations. BioGRID [Oughtred et al., 2021] and STRING [Szklarczyk et al., 2025] provide independent sources of positive interactions; their evaluations pair 10,111 source-specific positives with constructed negatives. Appendix Table S1 lists the evaluated sets and keeps assay non-detections, curated negatives, and constructed unobserved pairs explicitly separate.

These external sources are not assumed to be protein-or family-disjoint from Gold. Protein identities, homologous sequences, or previously observed pairs can therefore overlap across sources. The source-overlap analysis identifies, for example, 109 exact Y2H-iso sequences and four exact Y2H-iso sequence pairs overlapping Gold train; for STRING, raw-sequence matching finds 3,550 exact sequences and 33,571 exact sequence pairs overlapping Gold train. These counts describe overlap in the source pools; they do not imply that every evaluated protein pair overlaps Gold training data. We therefore use these datasets to test cross-source transfer without adaptation, not to claim strict OOD generalization. Appendix Figure S10 reports the corresponding overlap analysis.

### 3.2 PPI prediction models

We chose four models before final test inspection to span distinct sequence-only architectures while remaining feasible to retrain for six negative constructions and three seeds. Inclusion required a scalar pair score, training and inference behavior that could be validated, and no unavoidable PPI-specific pair pretraining under a fixed negative construction that would confound the controlled intervention. **Frozen-ESM2** uses fixed single-protein ESM-2 (150M) representations [Lin et al., 2023] and a symmetric pair MLP; the ESM-2 encoder is pretrained on protein sequences, but the interaction head is trained from scratch for each Gold training construction. **TUnA** uses ESM-2 per-token representations, intra-protein and inter-protein Transformer encoders, and an SNGP output layer [Ko et al., 2024]. **D-SCRIPT-ESM2** is the ESM-2-based (650M) D-SCRIPT implementation evaluated by Reim et al. [2025], derived from Sledzieski et al. [2021]. **RAPPPID** is a raw-sequence twin AWD-LSTM model with SentencePiece tokenization and no pretrained pLM [Szymborski and Emad, 2022]. Exact source-code revisions and architecture/training specifications are summarized in Appendix Table S2 and released with the experiment configuration.

### 3.3 Six negative constructions

We evaluate six protocols on the fixed Gold positive backbone. The published Gold benchmark used random negative sampling while approximately preserving protein degree; our Degree-matched construction is the closest controlled analogue but is regenerated under common candidate rules [Bernett et al., 2024, Joeres et al., 2026]. Figure 1 summarizes the construction and evaluation design.

**Figure 1:**
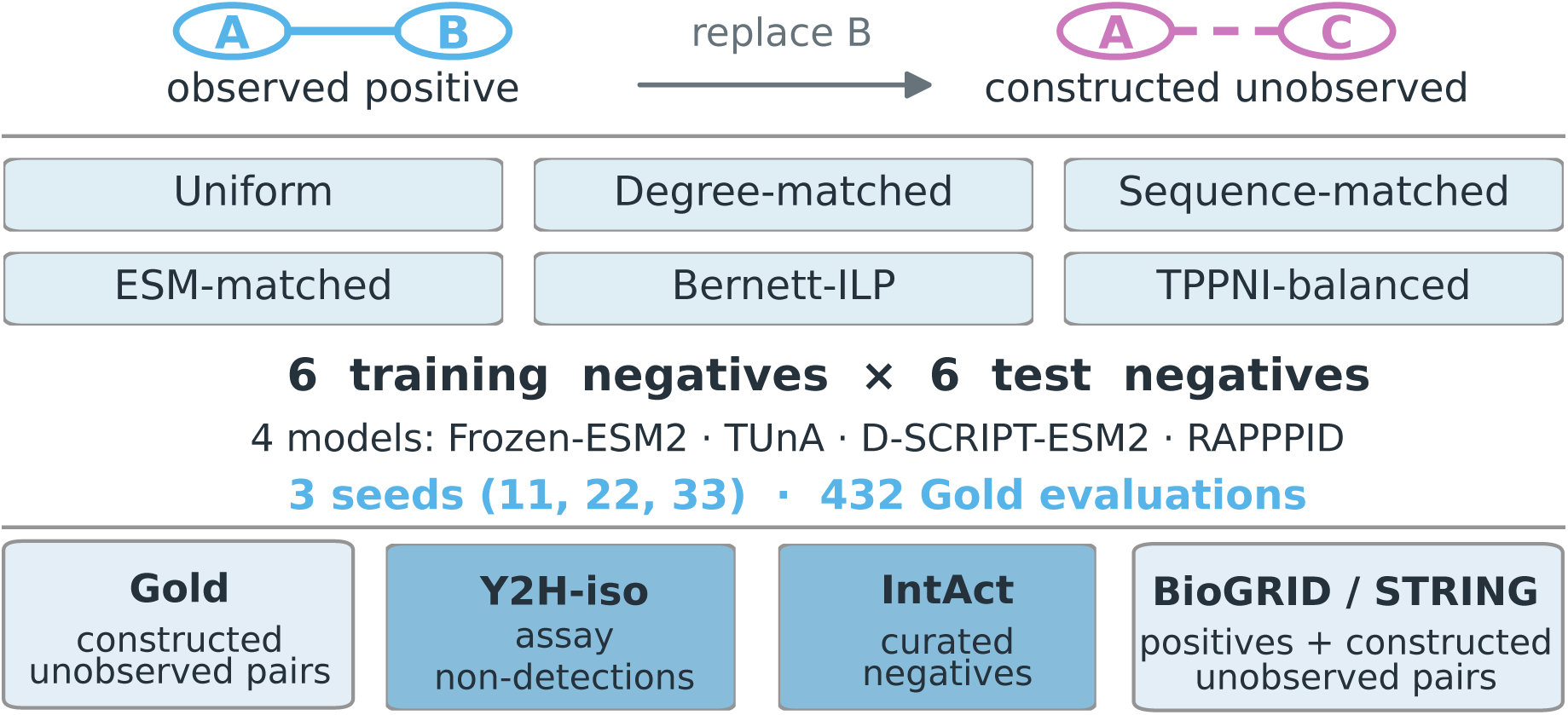
Negative-pair construction, Gold factorial, and evaluation evidence. Observed Gold positives are kept fixed while the partner used to construct a negative pair changes under one of six predeclared rules. The Gold experiment crosses six training-negative constructions, six test-negative constructions, four models, and three seeds, giving 432 evaluations. Cross-source evaluation keeps Y2H-iso assay non-detections, explicit IntAct curated negatives, and constructed BioGRID/STRING unobserved pairs distinct. Exact dataset counts are in Appendix Table S1.

#### Uniform

A replacement partner is sampled uniformly from the stage-specific candidate set. Uniform sampling does not match per-protein properties between classes and therefore leaves protein-level shortcuts available.

#### Degree-matched

The replacement partner minimizes the absolute difference from the train-visible network degree of the observed positive partner. On the protein-disjoint Gold test set, train-visible degree is degenerate because test proteins are absent from the training network, so this test construction is retained as a diagnostic condition.

#### Sequence-matched

Each protein is represented by log(1 + length) and the frequencies of the 20 standard amino acids, standardized within the split; the allowed replacement partner minimizing Euclidean distance to the observed partner is selected.

#### ESM-matched

The same fixed-anchor rule is applied in cached single-protein ESM-2 (150M) representation space, using final-layer overlap-corrected mean-pooled 640-dimensional vectors [Lin et al., 2023]. The representation cache is fixed before PPI training.

#### Bernett-ILP

We implement the global ILP negative construction of Bernett et al. [2026] on the fixed candidate-pair set, with equal weights on degree, taxon-pair counts, self-interactions, and mean Gene Ontology Biological Process Jaccard discrepancy.

#### TPPNI-balanced

We reproduce the degree-constrained configuration-model and Contrastive-L3 candidate generation of Chatterjee et al. [2025], then take a predefined deterministic 1:1 subset.

The six Gold test negative sets share nearly the same proteins (pairwise protein-set Jaccard 0.94–1.00) but few exact pairs (maximum off-diagonal Jaccard 0.048, Sequence-matched versus ESM-matched). Since each construction selects 10,111 pairs from a much larger candidate pool, low pair overlap does not by itself imply different distributions. As a descriptive measure of pair geometry (not of biological realism), the mean cosine similarity between the two ESM-2 representations in a pair is 0.908 for positives and 0.885–0.912 across negative sets, from TPPNI-balanced to ESM-matched. Full overlap statistics, similarity distributions, and per-partition overlap are given in Appendix Figures S7, S8, and S9.

For BioGRID and STRING we use a predeclared reduced set of only three constructions, spanning the random baseline (Uniform), the global bias-aware ILP construction (Bernett-ILP), and the topology-driven SCM/L3 construction (TPPNI-balanced), rather than duplicating the full six-construction Gold factorial on each external source.

## 4 Experimental Protocol and Controls

### Training-negative by test-negative evaluation

Each model is trained with each of the six Gold negative constructions using seeds 11, 22, and 33, giving 72 trained model instances. Every trained model is evaluated on all six fixed Gold test constructions, giving 432 evaluations. For a model trained on construction *c*, checkpoint selection uses maximum AUPRC on the matched Gold validation construction *c*; the classification threshold is selected by maximum MCC on that validation set. No test protein, test label, or external label is used for training, checkpoint selection, threshold selection, sampler fitting, or model inclusion.

All six main Gold test constructions are balanced 1:1. We report both AUPRC and AUROC as threshold-free classification metrics; AUPRC remains the pre-specified checkpoint-selection metric. The pre-specified evaluation protocol also records MCC, balanced accuracy, F1, Brier score, negative log-likelihood, and expected calibration error as secondary diagnostics; balanced accuracy, MCC, and F1 are shown in Appendix Figures S11, S12, and S13. Threshold-dependent metrics use the validation-selected threshold and are never optimized on test or external labels. External evaluation sets retain their native class prevalence; we use AUROC for cross-source discrimination comparisons and interpret AUPRC within each source relative to its prevalence baseline, rather than comparing raw AUPRC across sources with different class prevalence.

### Cross-source and leakage-reduced evaluation

Gold-trained models are evaluated without retraining or adaptation on Y2H-iso, IntAct, BioGRID, and STRING. We use *cross-source* only to denote a change of database or assay provenance; it does not imply sequence or family separation. Strict OOD claims are reserved for evaluations that additionally satisfy a declared protein-or family-separation rule. Separately, we evaluate D-SCRIPT-ESM2, Frozen-ESM2, and TUnA on the DataSAIL Gold-v4 sensitivity split [Joeres et al., 2026]. DataSAIL reduces the reported scaled leakage score for Gold from 0.3642 to 0.0465, but the published Gold-v4 split also changes which interactions are retained. We therefore report it as a separate leakage-control analysis rather than a matched replacement for the main six-construction experiment.

### Controls that do not require the original protein pair

The **easy-property** classifier uses predefined dataset properties such as protein degree, sequence length, and amino-acid composition. The **single-protein sequence** classifier represents one endpoint at a time by length and amino-acid composition and averages endpoint probabilities. The **simple pooled-ESM** classifier mean-pools final-layer ESM-2 residue embeddings and fits a low-capacity logistic classifier to *u* + *v* and *|u − v|*. Finally, **partner-shuffled evaluation** keeps each trained PPI model fixed but deterministically permutes the second protein within each label class, preserving label counts and single-protein marginals while breaking the original pairs. We define Δ_pair_ = AUROC_original_ *−* AUROC_shuffled_; positive values favor the original pairing and negative values favor shuffled partners.

The exact seeds, source-code revisions, candidate-pair definitions, negative-set hashes, validation rules, model configurations, and machine-readable figure data are released with the experiment configuration.

## 5 Results

### Performance depends on both training and test negative construction

With Uniform negatives for both training and testing, mean AUPRC is 0.723 for TUnA, 0.697 for Frozen-ESM2, 0.587 for D-SCRIPT-ESM2, and 0.518 for RAPPPID. These values are not stable when the negative construction changes (Fig. 2 and Appendix Figure S1). Holding the trained model fixed and changing only test negatives gives an average AUPRC range of 0.165 for TUnA and 0.147 for Frozen-ESM2, with maximum ranges of 0.249 and 0.264. Holding the test set fixed and changing only training negatives gives average ranges of 0.138 and 0.109, respectively. Moreover, negative construction influences model ordering. Across the 36 train-by-test settings, Frozen-ESM2 has the highest mean AUPRC in 20, TUnA in 15, D-SCRIPT-ESM2 in one, and RAPPPID in none. The separate reversal analysis finds 290 eligible model-pair ordering changes, of which 178 are also supported by paired evaluation resampling and seed-to-seed consistency. These counts summarize the tested settings and do not define a universal model ranking. Matching the training and test negative construction does not consistently improve performance (Appendix Fig. S2).

**Figure 2:**
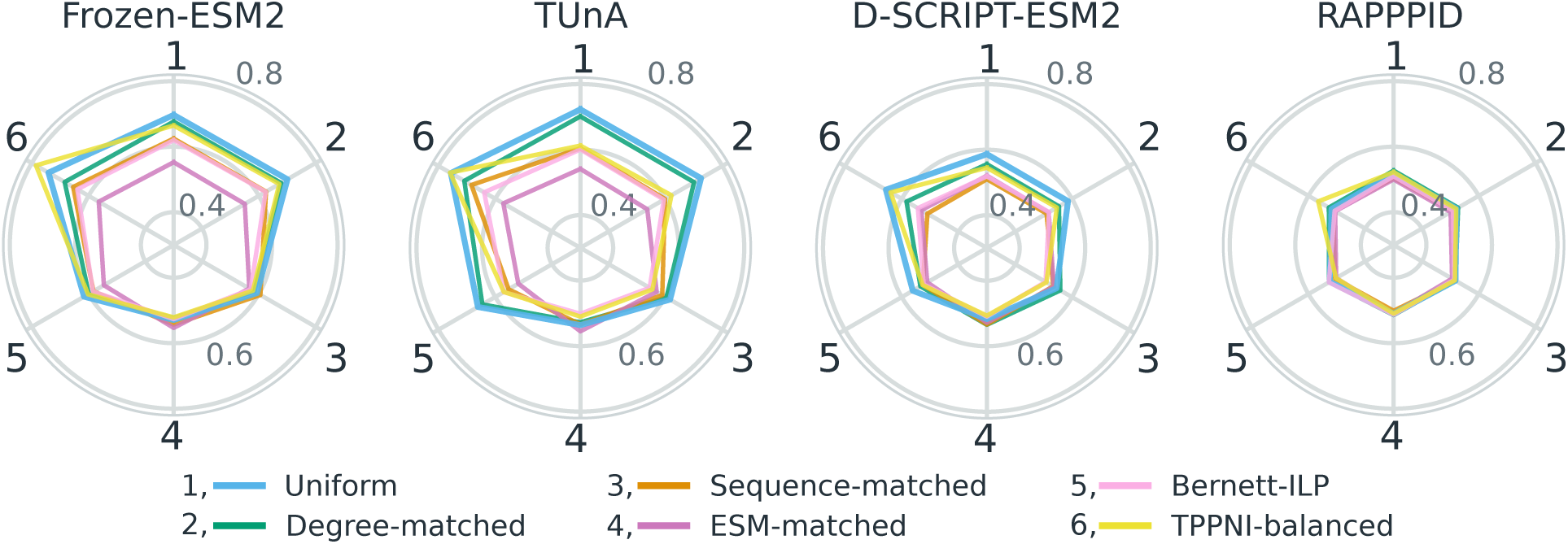
Gold training-by-test factorial. Four radar panels show absolute mean AUPRC over seeds 11, 22, and 33. Spokes 1–6 are the Gold test-negative constructions (Uniform, Degree-matched, Sequence-matched, ESM-matched, Bernett-ILP, and TPPNI-balanced), and colored lines are the training-negative constructions. Exact train-by-test values are given in Appendix Figure S1.

### Different negative constructions leave different shortcuts available

Reducing one shortcut does not remove all simple predictive signals. Across the six Gold test constructions, the AUROC of the easy-property, single-protein sequence, and simple pooled-ESM controls changes substantially, and the strongest diagnostic is not the same in every negative construction (Appendix Figure S3). These controls are not intended as competing PPI predictors or as state-of-the-art baselines; they are diagnostic probes asking whether labels remain predictable without modeling the specific compatibility of the original protein pair. We therefore use them to identify which simple signals remain available under each construction, rather than to rank the controls against the four PPI models.

### Partner shuffling tests dependence on the original pair

Within-class partner shuffling tests whether a model needs the original A–B pairing to retain its AUROC; Δ_pair_ *>* 0 indicates that it does. Across models and test sets, Δ_pair_ never exceeds 0.012, reached on TPPNI-balanced by Frozen-ESM2 and TUnA (Fig. 3 and Appendix Figure S4). It is small and mixed for D-SCRIPT-ESM2, and close to zero for RAPPPID, as expected given its near-chance performance. For Frozen-ESM2 and TUnA, Δ_pair_ is negative on Sequence-matched (*−*0.047, *−*0.050), ESM-matched (*−*0.086, *−*0.087), and, less so, Bernett-ILP (*−*0.029, *−*0.023): these models score higher on shuffled pairs than on the original ones. Sequence-and ESM-matched negatives are anchor-based: each positive (*A, B*) has a negative (*A, C*) with *C* nearest to *B* in the matching space, so the two pairs share *A* and have near-identical partners and receive similar scores. Shuffling preserves the protein composition of each class but breaks this one-to-one correspondence, which makes the classes easier to separate. Bernett-ILP matches class-level statistics rather than individual pairs, and the effect is correspondingly smaller. In no construction does the original pairing contribute substantially to AUROC.

**Figure 3:**
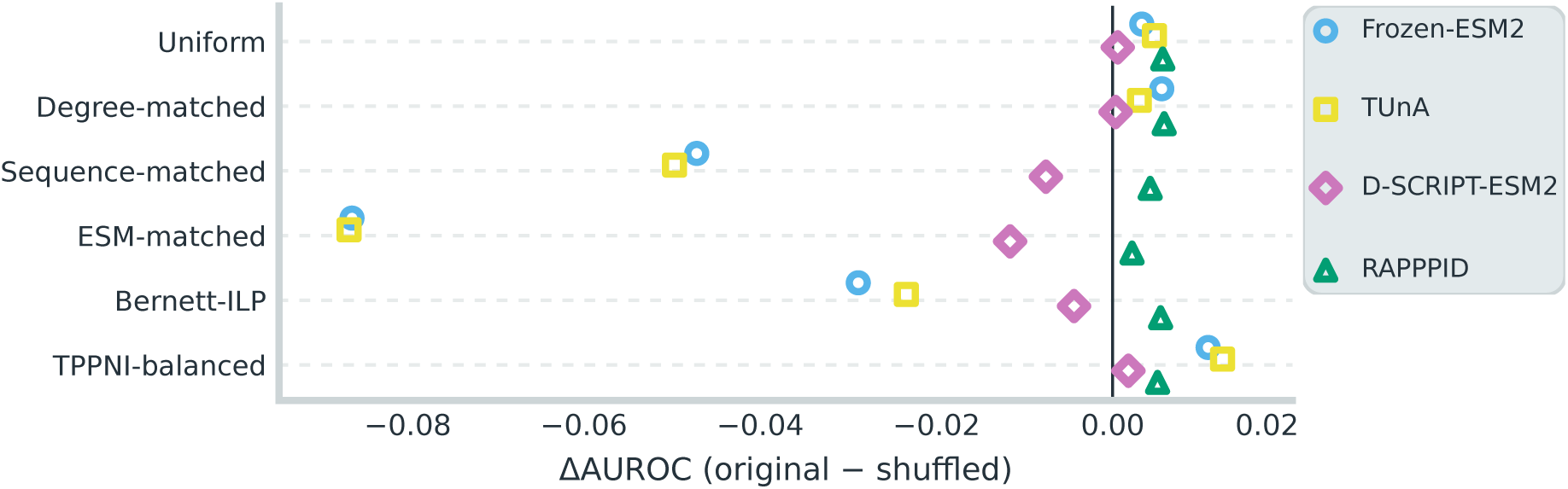
Effect of breaking the original protein pairs. Points show ΔAUROC = AUROC(original pairs) *−* AUROC(shuffled pairs) for each model and Gold test-negative construction, averaged over seeds 11, 22, and 33. Positive values favor the original pairing; negative values favor shuffled partners. The vertical zero line marks no change. Absolute AUROC values are shown in Appendix Figure S4.

### Constructed training negatives rarely improve external transfer

We compare each alternative training construction with Uniform training on the same external evaluation set. For the three ESM2-based models, constructed training negatives underperform Uniform training in 26 of 30 comparisons on sources with assay-based or curated negatives (Y2H-iso, IntAct) and in 82 of 90 on BioGRID and STRING with constructed negatives (Appendix Figure S5), with losses of up to 0.229 AUROC. Training on the construction used to build an external set does not help either: TPPNI-balanced training underperforms Uniform on both TPPNI-constructed external sets for all three models (STRING and BioGRID, Appendix Figure S5). Gains are small and specific. Degree-matched training improves transfer to Bernett-constructed external sets (BioGRID, STRING) for Frozen-ESM2 and TUnA (at most +0.051), consistent with the ILP matching protein degree. On sources with assay non-detections and curated negatives, TPPNI-balanced training slightly improves Y2H-iso AUROC for all three ESM2-based models (at most +0.037), and Bernett-ILP training improves IntAct AUROC for D-SCRIPT-ESM2 (+0.028). Some gains coincide with lower Gold performance, for example TPPNI-balanced training for Frozen-ESM2 (Gold AUPRC 0.697 to 0.665, Y2H-iso AUROC 0.598 to 0.620; Fig. 4). A construction can thus trade constructed-benchmark performance for external transfer, but such trade-offs are rare, and making negatives resemble positives helps only in a limited sense: matching protein degree costs little relative to Uniform and helps on external sets built with degree matching, whereas matching individual partners (Sequence-and ESM-matched) causes the largest external losses.

**Figure 4:**
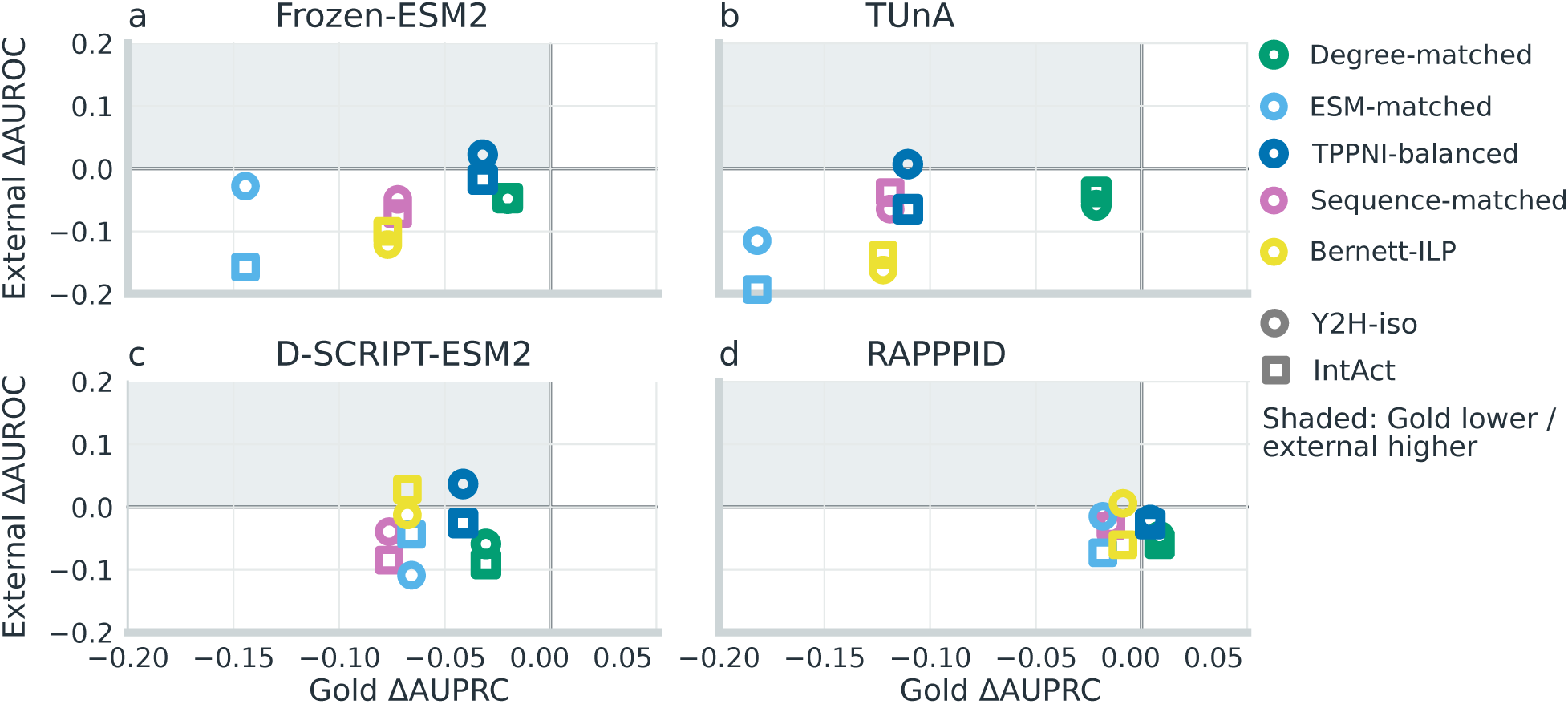
Gold benchmark change versus cross-source transfer to assay-based and curated negatives. Each panel corresponds to one model, and each point to one alternative training construction, compared with Uniform training of the same architecture. The x-axis is ΔAUPRC on the fixed Uniform Gold test; the y-axis is ΔAUROC on Y2H-iso (circles) or IntAct (squares), as in Appendix Figure S5. The shaded upper-left quadrant marks lower Gold AUPRC together with higher measured-source AUROC. Absolute Gold AUPRC values are in Appendix Figure S1.

### DataSAIL provides a separate leakage-control analysis

The published DataSAIL Gold-v4 sensitivity analysis uses Uniform and ESM-matched negatives for both training and testing and includes three of the four models. Mean AUPRC across its tested conditions is 0.529 for D-SCRIPT-ESM2, 0.582 for Frozen-ESM2, and 0.584 for TUnA. Because this analysis changes both the split and sequence subset and covers only two negative constructions, it remains a separate sensitivity analysis rather than a replacement for the main factorial (Appendix Fig. S6).

## 6 Discussion and conclusion

Dataset-level shortcuts, pair dependence, benchmark difficulty, and cross-source transfer are distinct properties of a PPI benchmark and require separate measurements rather than a single score. High performance on a balanced constructed test set shows that a model separates observed positives from the negatives defined by that benchmark; it does not establish performance on arbitrary protein pairs or experimentally established non-interactions.

Negatives matched in ESM-2 space are close to chance for all three ESM2-based models, whatever the training construction, while topology-driven negatives are the easiest. Partner shuffling shows where this difficulty lies: on anchor-matched test sets, breaking the pair correspondence increases AUROC for Frozen-ESM2 and TUnA, thus the difficulty comes from how negatives are placed relative to positives rather than from pair-specific signal the models fail to use. A benchmark’s difficulty should therefore be reported together with the space in which its negatives were matched.

Constructed training negatives rarely outperform uniform training outside their own construction, and anchor-matched training brings little benefit even on its own test set; models trained on ESM-matched negatives remain close to chance everywhere. Nearest-neighbour matching may reduce the diversity of the negative class, and may introduce unlabelled true interactions, since close homologs of a partner often share its interactions; in anchor-based constructions, the same anchor then appears with near-identical partners under opposite labels. In addition, reducing a known shortcut does not guarantee the best assay-based performance: Bernett-ILP removes the easy-property signal but is not best on Y2H-iso or IntAct, whereas TPPNI-balanced training improves Y2H-iso AUROC for two models despite an easily separable test construction. No single diagnostic makes a construction globally “realistic” or “unbiased”. Conversely, the frozen-embedding MLP is most often the best model, suggesting that pretrained representations carry most of the usable signal in this setting.

Our experiments complement Bernett et al. [2026], which reduces known positive/negative differences; PPIShift asks how conclusions change across constructions and evaluation sources. Negative construction should be treated as an evaluation axis rather than a preprocessing detail: report the candidate set and negative evidence; do not assume that harder training negatives improve transfer; test at least one alternative construction, including one matched in the model’s own representation space; include controls that do not require the original pair; and evaluate on external assay and curated data when possible, keeping cross-source transfer distinct from protein-or family-separated OOD generalization.

## 7 Limitations

The common 512-residue subset retains only 16.3% of the original Gold training positives (Appendix Fig. S14); results may not hold for the full Gold release or longer proteins, and the near-chance performance of RAPPPID may reflect this reduced training set. We test four models and a single pLM family, and pretrained representations may have been exposed to downstream proteins despite the protein-disjoint split [Szymborski and Emad, 2026]. None of our negative labels, whether constructed, assay-based, or curated, proves that two proteins cannot interact. The shortcut controls cover only selected properties, and partner shuffling tests whether the original pair is needed, not which mechanism a model uses. Three seeds quantify optimization variability, not biological replication.

## 8 Reproducibility statement

We provide an anonymized implementation of the dataset-processing pipeline, negative constructions, model implementations, experiment configurations, evaluation scripts, and source data used to generate the figures at https://github.com/hanzopgp/PPIShiftSubmission. Released files record random seeds, evaluation-set and checkpoint hashes, and validation-only selection rules. Machine-readable main, external, DataSAIL, and control results are included together with the data underlying every main figure.

## 9 Acknowledgments

This work was granted access to the HPC resources of the LIP6 Convergence cluster at Sorbonne Université. This work was supported by a doctoral grant from the French Ministry of Higher Education and Research (MESR). It has been co-funded by the European Union (ERC, PROMISE, 101087830). Views and opinions expressed are however those of the author(s) only and do not necessarily reflect those of the European Union or the European Research Council. Neither the European Union nor the granting authority can be held responsible for them. For the purpose of open access, a CC-BY public copyright licence has been applied by the authors to the present document and will be applied to all subsequent versions up to the Author Accepted Manuscript arising from this submission. This work has also been co-funded by the French National Agency for Research (ANR, France 2030, PostGenAI@Paris, ANR-23-IACL-0007).

## **A** Dataset and Model Details

### Frozen-ESM2

The fixed encoder is ESM-2 (150M) [Lin et al., 2023]. Residue embeddings from the final (30th) layer are overlap-corrected and mean-pooled to one 640-dimensional vector per protein. The three symmetric feature blocks are standardized on the training set before the MLP. The head uses ReLU activations and Adam with learning rate 10*^−^*^3^ and weight-decay/regularization coefficient 10*^−^*^4^ for at most 20 epochs; the epoch with maximum matched-validation AUPRC is kept.

### D-SCRIPT-ESM2

The implementation follows the D-SCRIPT contact-map architecture [Sledzieski et al., 2021, Reim et al., 2025] with frozen ESM-2 (650M) residue representations. Each residue is projected from 640 to 100 dimensions with ReLU and dropout 0.5. For every residue pair, absolute difference and element-wise product are concatenated, followed by a 200*→*50 1 *×* 1 convolution, batch normalization, a 50*→*1 7 *×* 7 contact convolution, and the D-SCRIPT pooling parameters. The downstream model has 76,706 trainable parameters and is optimized for 10 epochs with Adam at 10*^−^*^3^ using binary cross-entropy; checkpoint selection uses matched-validation AUPRC.

### TunA

We use the fixed TUnA configuration [Ko et al., 2024]: a 640*→*64 projection, one intra-protein and one inter-protein Transformer layer, eight attention heads, feed-forward width 256, dropout 0.2, Swish activation, and an SNGP output layer with 4,096 random Fourier features. This gives 145,088 trainable downstream parameters with the ESM-2 encoder frozen. Training uses Adam+Lookahead with learning rate 10*^−^*^4^, weight decay 10*^−^*^5^, batch size 16, and 14 epochs; the retained checkpoint maximizes matched-validation AUPRC. Pair scoring follows the fixed swap-invariant inference rule used in this evaluation.

### RAPPPID

We use the RAPPPID architecture [Szymborski and Emad, 2022]: a training-only SentencePiece unigram vocabulary of 250, 64-dimensional token embeddings, a shared two-layer bidirectional AWD-LSTM with 32 hidden units per direction, embedding dropout 0.3, recurrent weight DropConnect 0.3, and a 128*→*32*→*1 Mish classifier with pair-head dropout 0.2. This configuration has 74,497 trainable parameters and no pretrained pLM. Training uses Ranger21 with stochastic weight averaging. Checkpoint selection uses matched-validation AUPRC under the common evaluation protocol; pair scores are made swap-invariant by the fixed inference rule used in this study.

Across all four models, one training unit is one model *×* training-negative construction *×* seed. The selected checkpoint is reused unchanged across all six test constructions. Exact source-code revisions, negative-set hashes, and serialized model configurations are included with the released experiment files.

**Table S1:** Evaluation datasets and the evidence supporting their negative labels. Assay outcomes, curated negatives, and constructed unobserved pairs are kept separate throughout the analysis.

| Dataset | Pos. | Neg. | Negative evidence |
| --- | --- | --- | --- |
| Y2H-iso [Lambourne et al., 2026] | 846 | 3,185 | assay non-detection |
| IntAct [Orchard et al., 2014] | 10,111 | 236 | explicit curated negative |
| Gold train [Bernett et al., 2024] | 13,304 | 13,304 | constructed pair, six constructions |
| Gold validation [Bernett et al., 2024] | 11,085 | 11,085 | constructed pair, six constructions |
| Gold test [Bernett et al., 2024] | 10,111 | 10,111 | constructed pair, six constructions |
| BioGRID [Oughtred et al., 2021] | 10,111 | 10,111 | constructed pair, three constructions |
| STRING [Szklarczyk et al., 2025] | 10,111 | 10,111 | constructed pair, three constructions |

## **B** Additional Results

### Execution summary

The final protocol completed all main planned evaluations: four models, six training-negative constructions, six test-negative constructions, and three seeds. External datasets were scored without retraining. The published DataSAIL Gold-v4 analysis is reported separately from the main Gold experiment.

### IntAct prevalence

The IntAct set contains 10,111 positives and 236 explicit negatives. Because positives dominate this set, its raw AUPRC is not compared with the 1:1 Gold AUPRC. The supplementary result tables report AUROC, AUPRC, MCC, balanced accuracy, and F1 for the native-prevalence evaluation set using the validation-selected threshold where required. We also report a fixed balanced subset as a post-hoc sensitivity check, kept separate from the native-prevalence result.

**Table S2:**
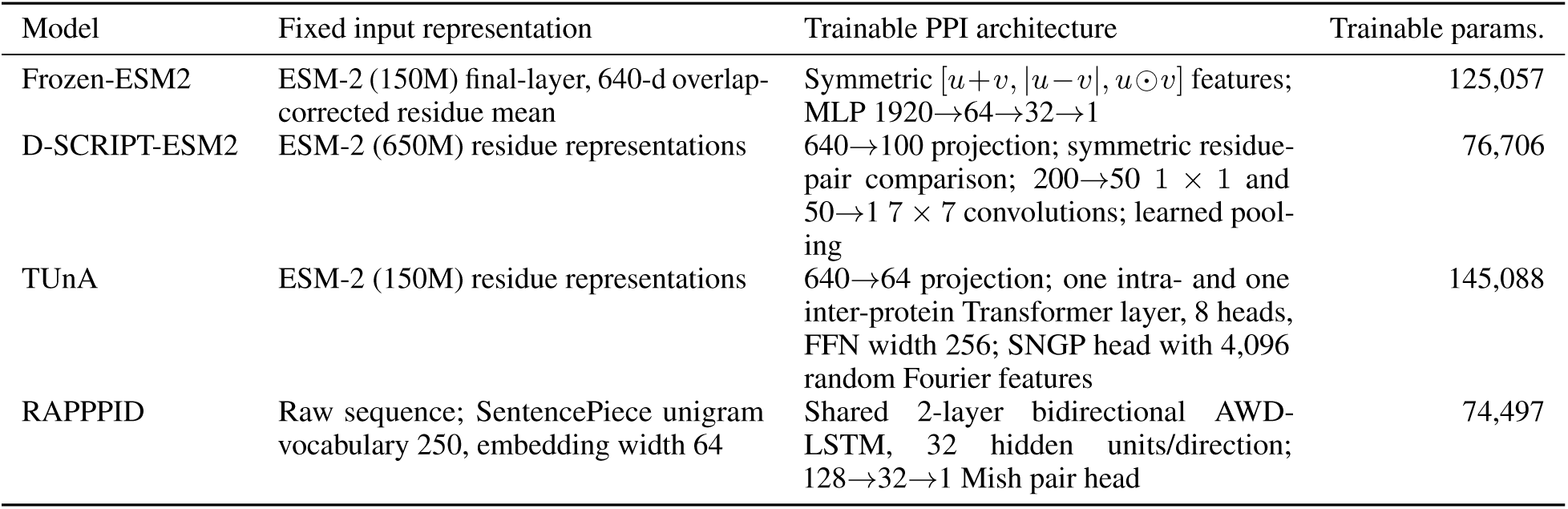
Core model architectures under the fixed model configurations. ESM-2 (150M) has 30 layers, approximately 150M parameters, and 640-dimensional residue outputs; it is frozen wherever used. Parameter counts below refer only to the trainable PPI model.

### Model-order reversals

The reversal analysis separates changes in point estimates from changes also supported by paired evaluation resampling and seed-to-seed consistency. The main text reports both counts; percentages are omitted here to avoid conflating different eligibility filters.

### DataSAIL

The published Gold-v4 split uses its own dataset definition and sequence-compatibility filter. RAPPPID was not run in this analysis and is not imputed.

## **C** Released Resources

The anonymous supplementary package contains the machine-readable results used for the main factorial experiment, external evaluations, DataSAIL sensitivity analysis, control experiments, calibration analyses, and reported figures. Configuration files record dataset provenance, negative-construction settings, random seeds, model configurations, and checkpoint identifiers needed to trace the reported analyses.

Additional released results include full training-by-test matrices, seed-level and bootstrap summaries, endpoint-frequency and selected-partner statistics, duplicate-pair sensitivity analyses, the pair-representation projection, and compute and hyperparameter records.

**Figure S1:**
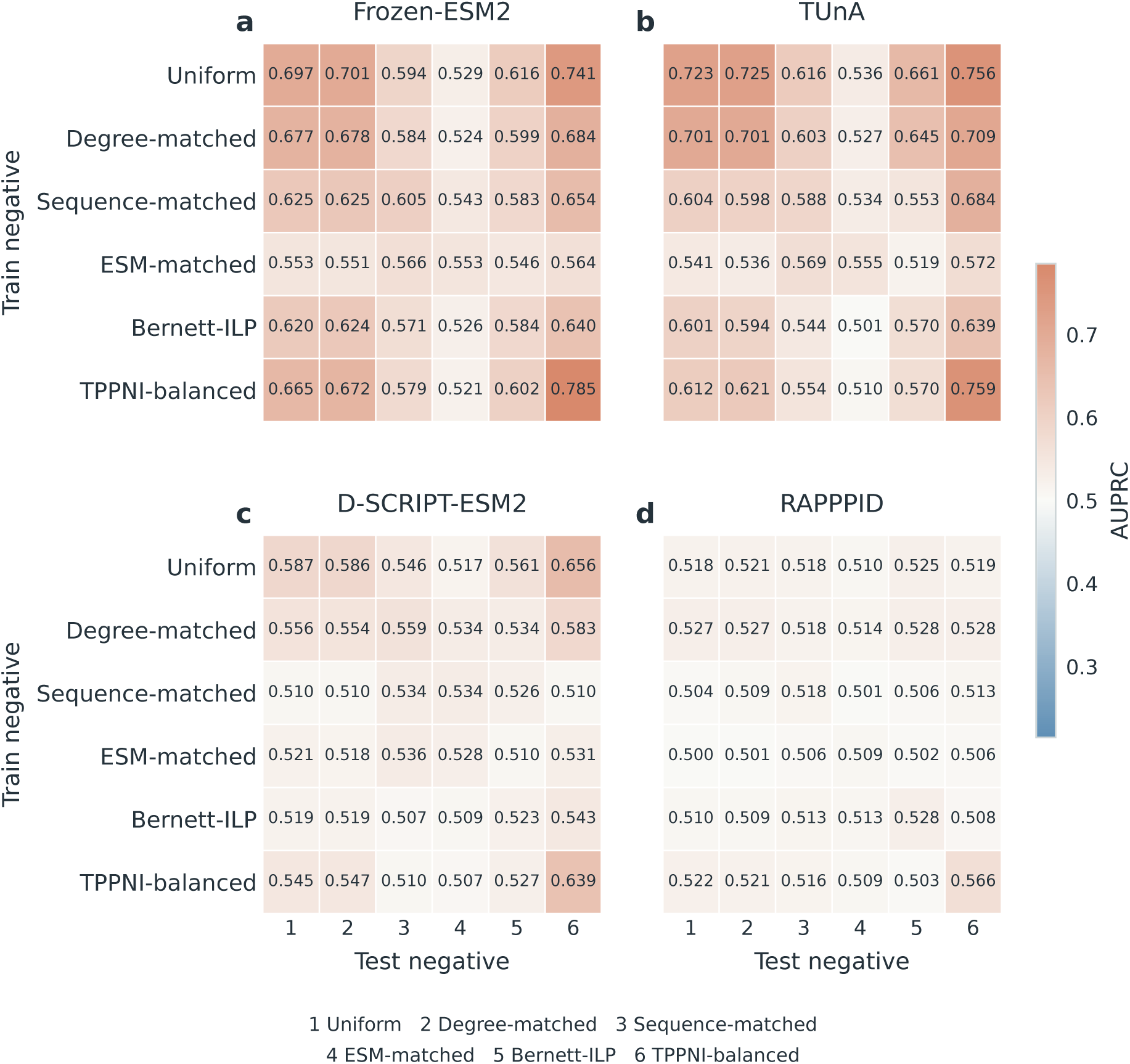
Cross-source transfer across Gold training-negative constructions. Each cell shows ΔAUROC = AUROC(trained on *c*) *−* AUROC(trained on Uniform) for the same architecture and external evaluation set, averaged over seeds; the Uniform column is zero by definition. Y2H-iso, IntAct, BioGRID, and STRING remain distinct evaluation sources and are not interpreted as strict OOD tests.

**Figure S2:**
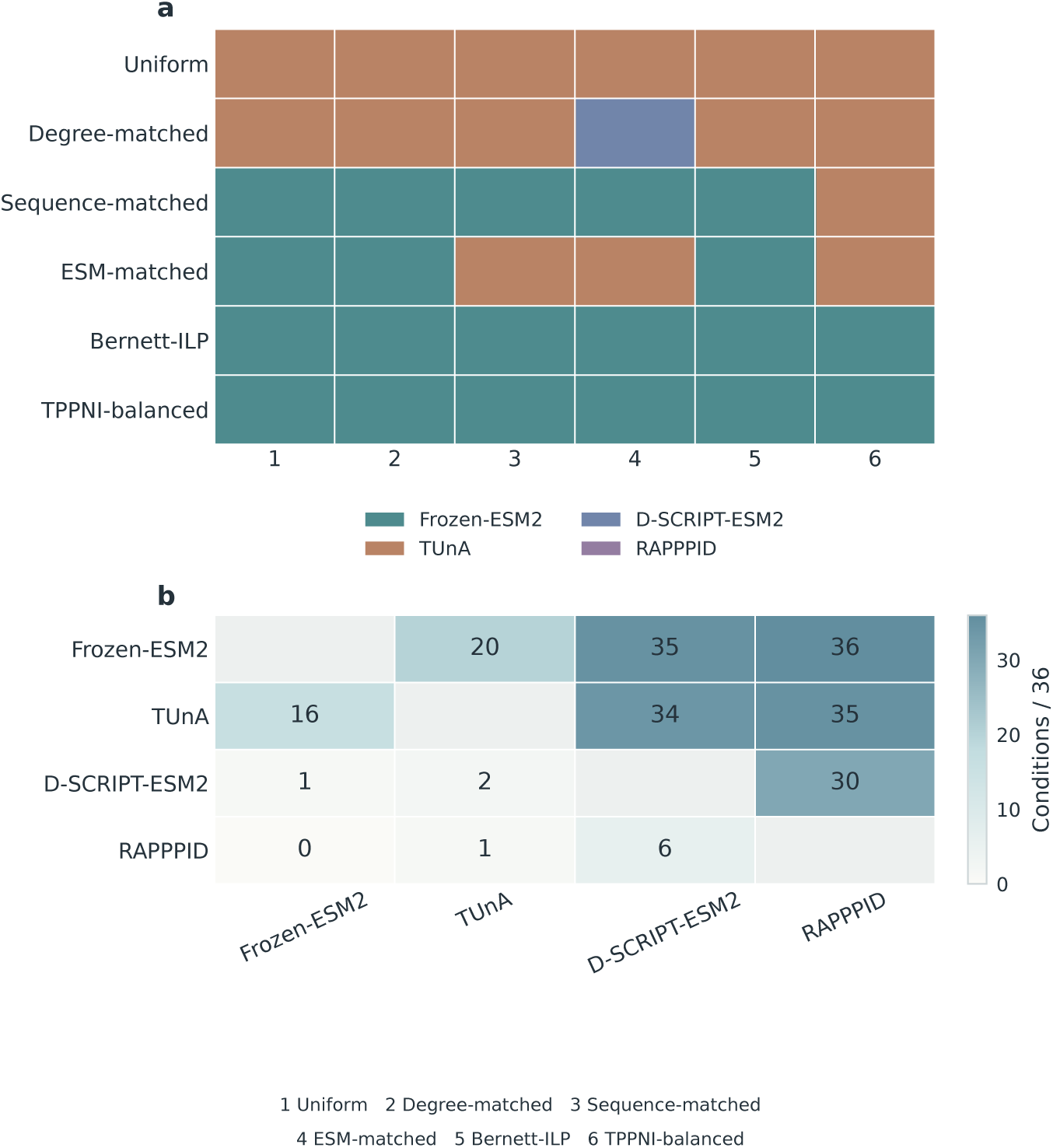
Model ordering depends on the negative construction. Panel a identifies the model with the highest mean AUPRC in each of the 36 tested train-by-test conditions. Panel b counts pairwise conditions in which the row model has higher mean AUPRC than the column model. These counts summarize the evaluated conditions and do not define a universal model ranking.

**Figure S3:**
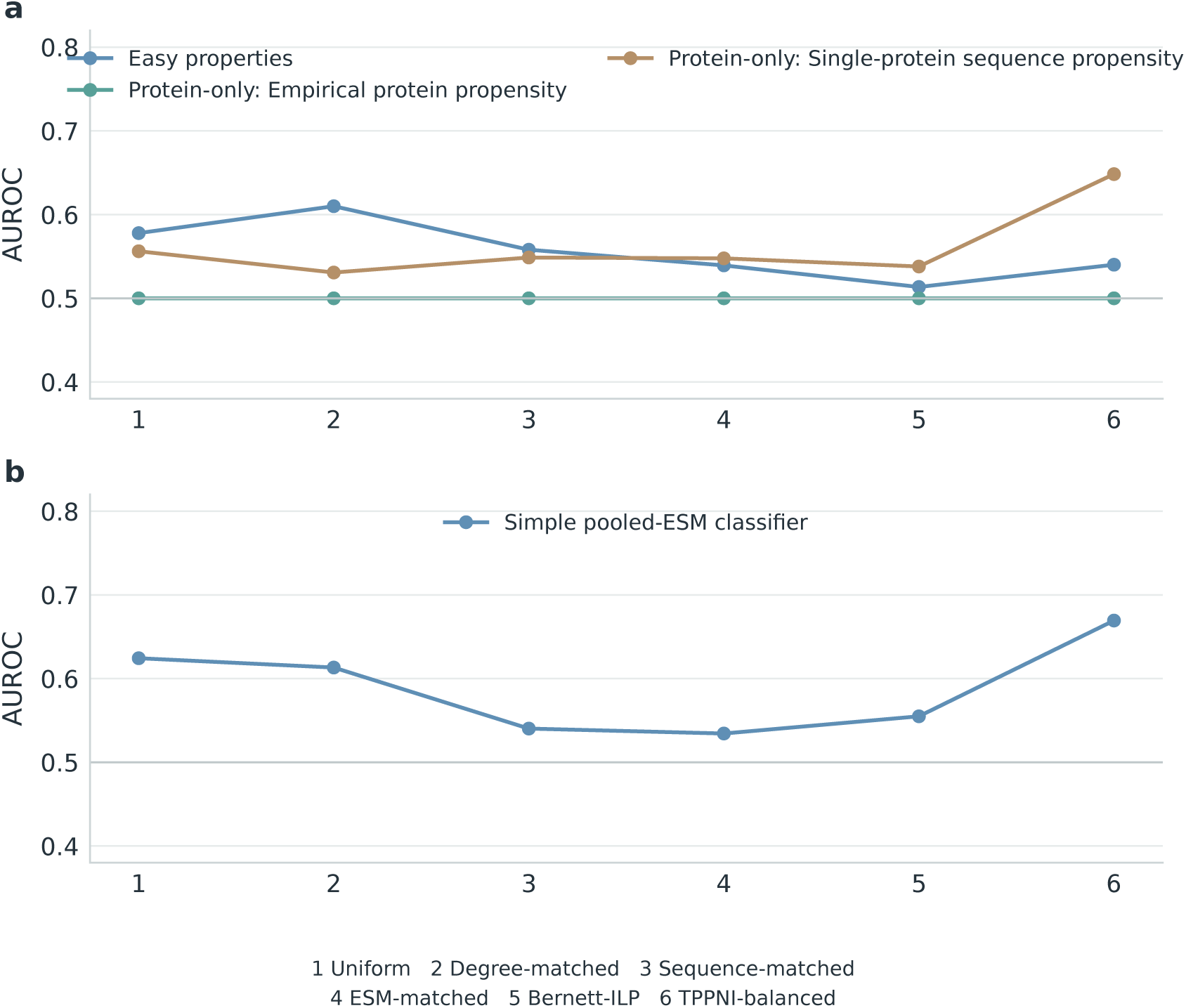
Additional shortcut-control diagnostics. Panel a shows protein-only and easy-property controls that do not require modeling the original protein pair; panel b shows the simple pooled-ESM classifier. The horizontal reference line marks AUROC = 0.5.

**Figure S4:**
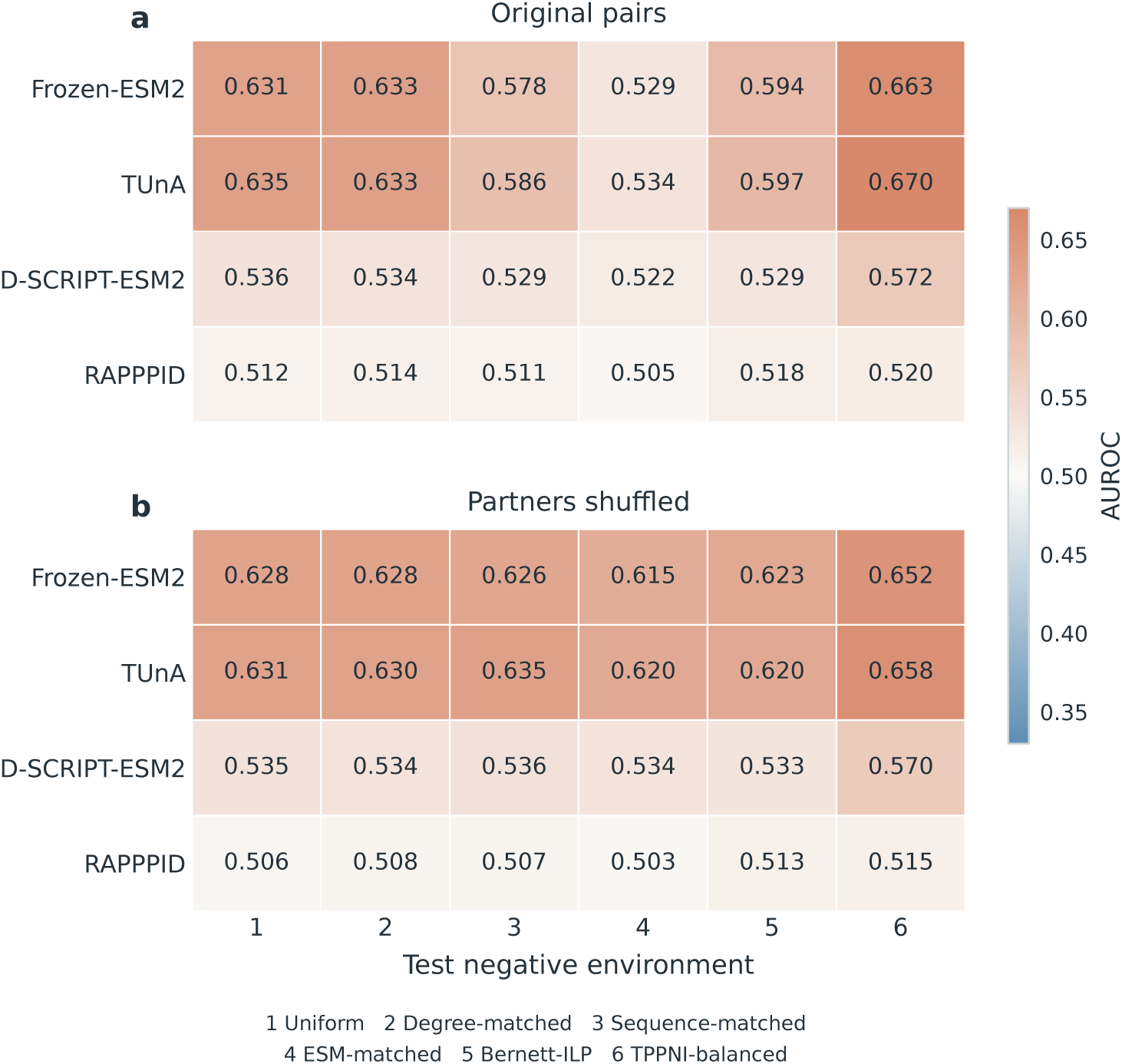
Absolute AUROC before and after partner shuffling. The same fixed checkpoints are evaluated on the original Gold test pairs and on the corresponding within-label partner-shuffled sets; no model is retrained after shuffling.

**Figure S5:**
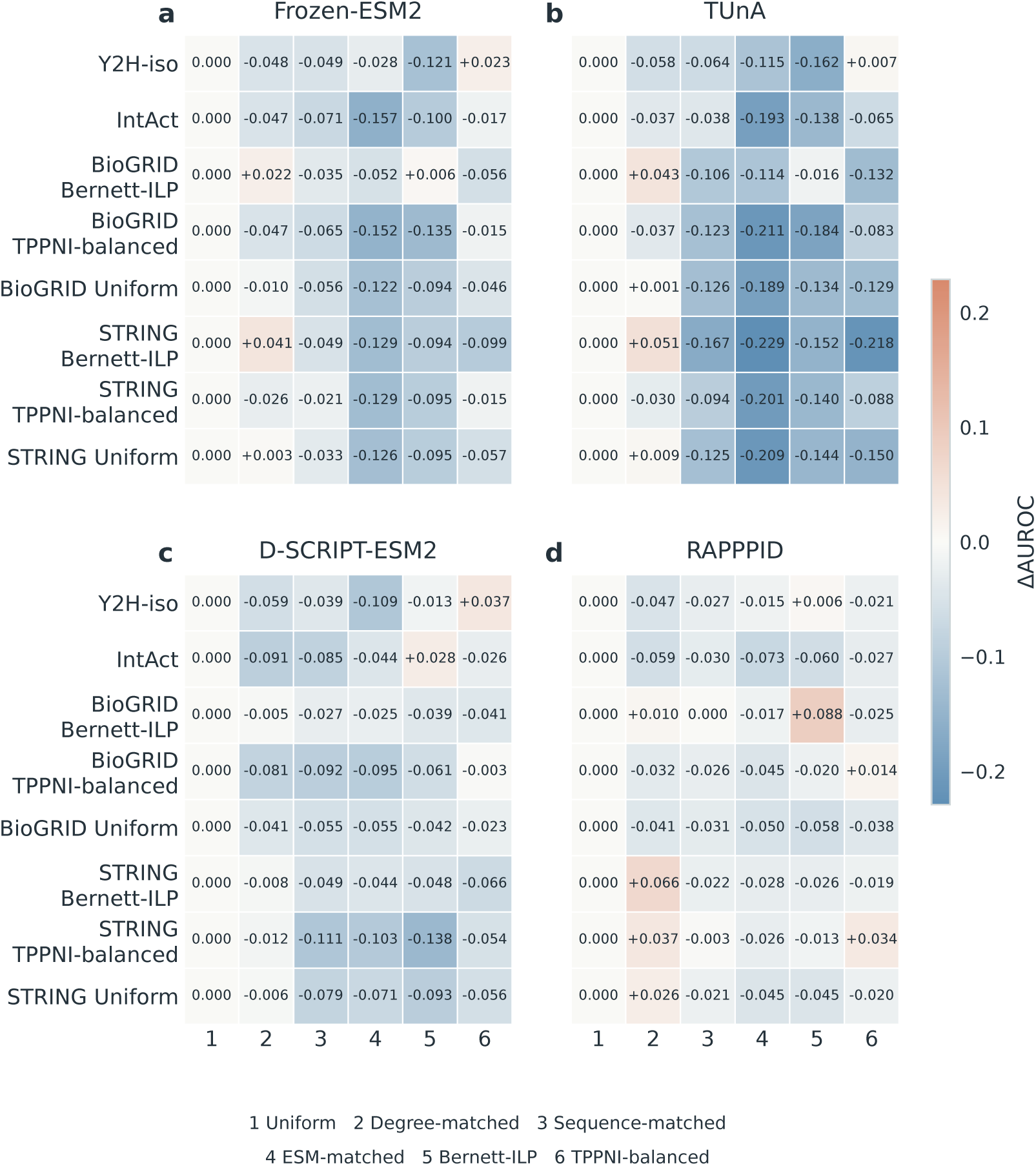
Cross-source transfer across Gold training-negative constructions. Each cell shows ΔAUROC relative to the corresponding Uniform-trained model on the same external evaluation set. Y2H-iso, IntAct, BioGRID, and STRING remain distinct evaluation sources and are not interpreted as uniformly strict OOD tests.

**Figure S6:**
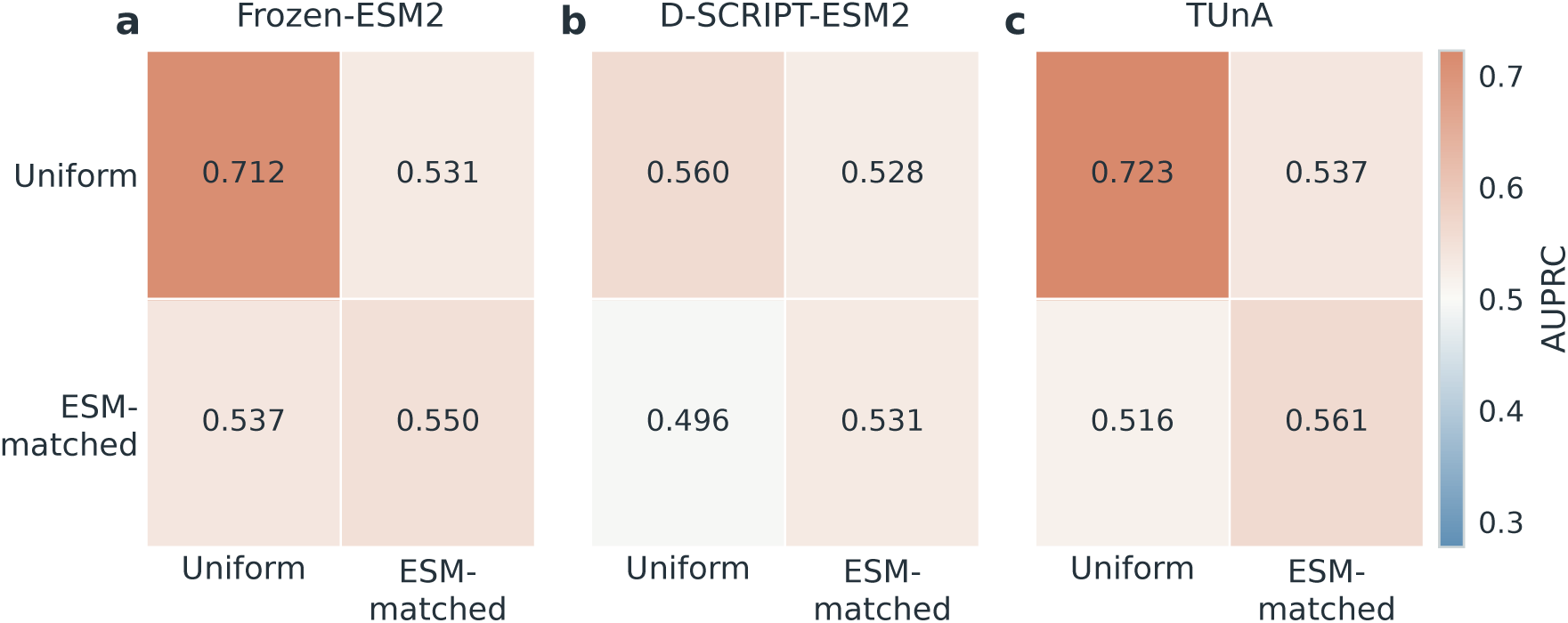
DataSAIL Gold-v4 sensitivity analysis. Mean AUPRC is shown for Uniform and ESM-matched training and test negative constructions for the three models evaluated on the published DataSAIL split. RAPPPID was not run in this analysis and is not imputed.

**Figure S7:**
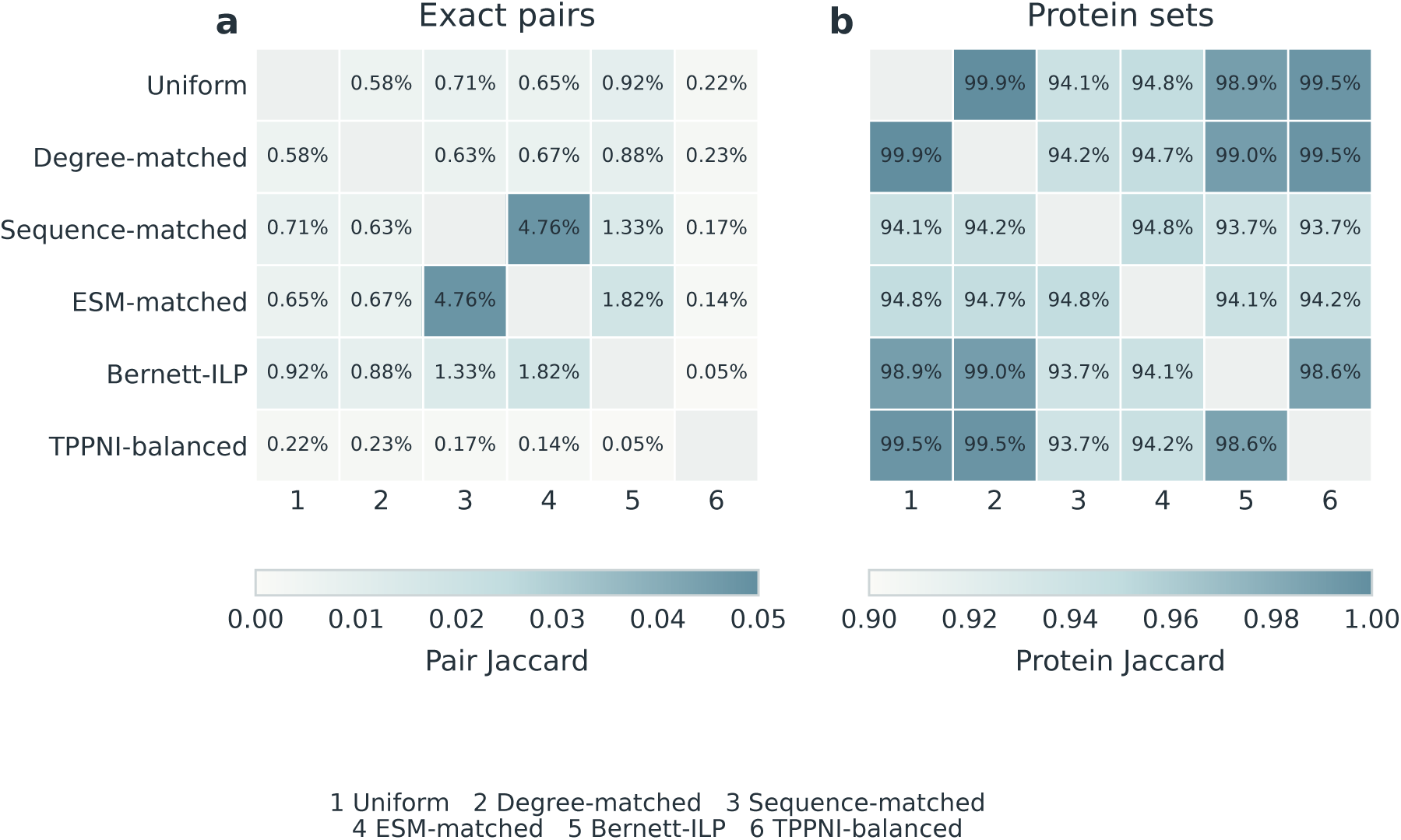
Overlap among the six Gold test negative constructions. Panel a shows exact-pair Jaccard overlap and panel b shows protein-set Jaccard overlap. Exact negative-pair overlap is low despite substantial reuse of the same proteins.

**Figure S8:**
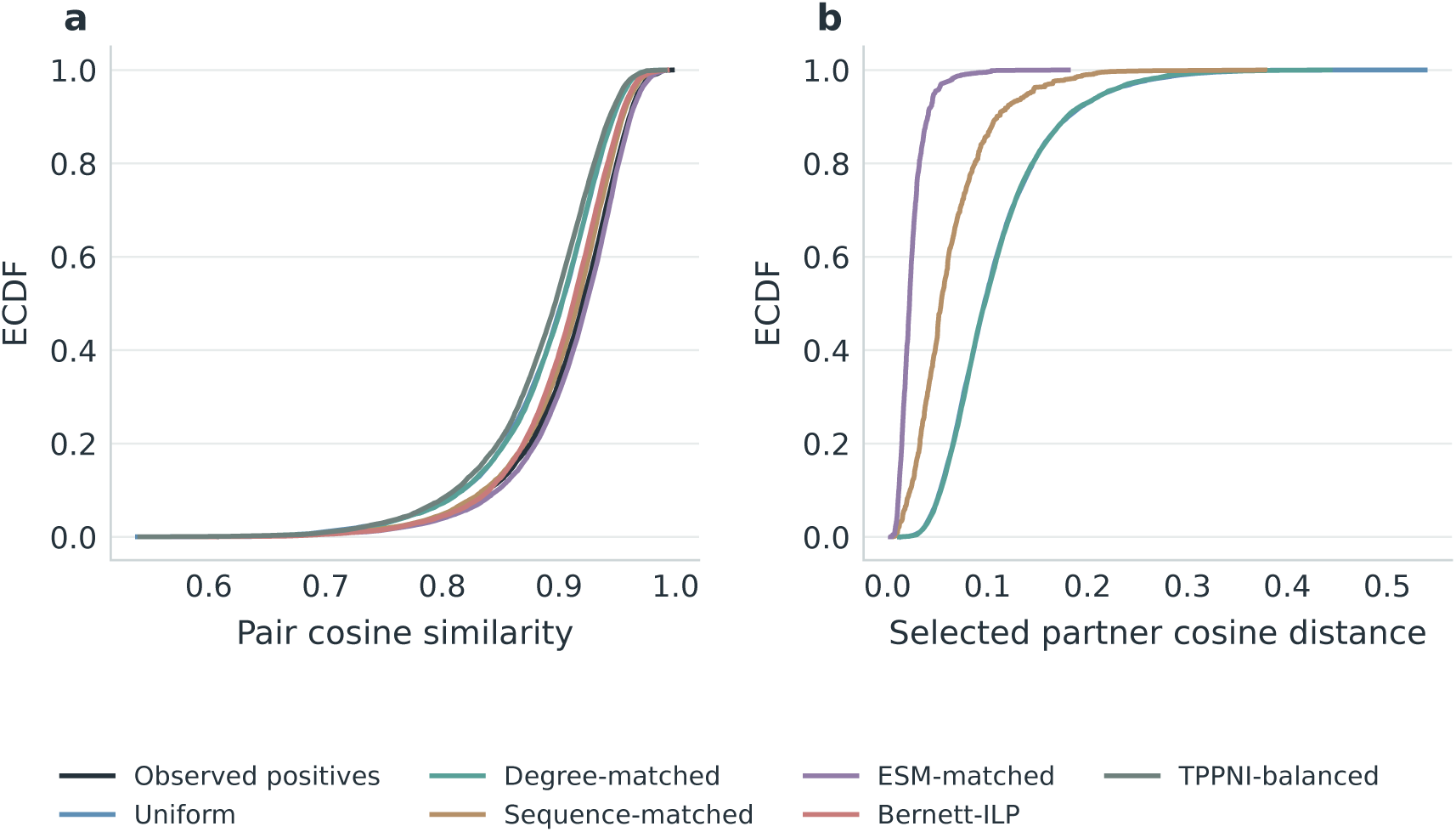
Representation-space characterization of the negative constructions. Panel a shows empirical distributions of within-pair ESM-2 cosine similarity. Panel b shows the cosine distance between the observed positive partner and the selected negative partner for constructions with a valid anchor-partner interpretation. These diagnostics are descriptive and do not rank biological realism.

**Figure S9:**
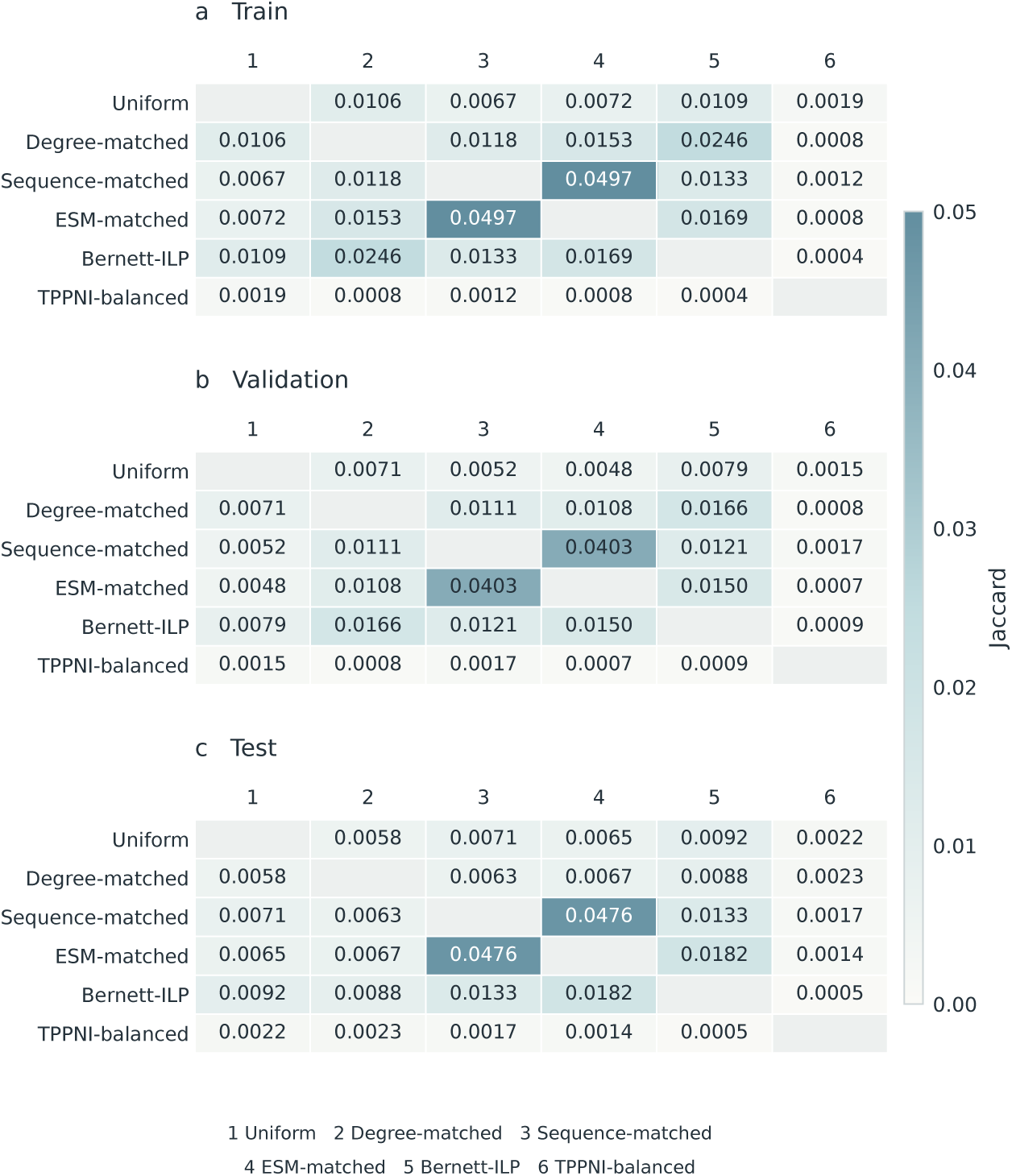
Negative-pair overlap across dataset partitions. Pairwise exact-pair Jaccard similarity is shown separately for the training, validation, and test negative constructions using a common scale.

**Figure S10:**
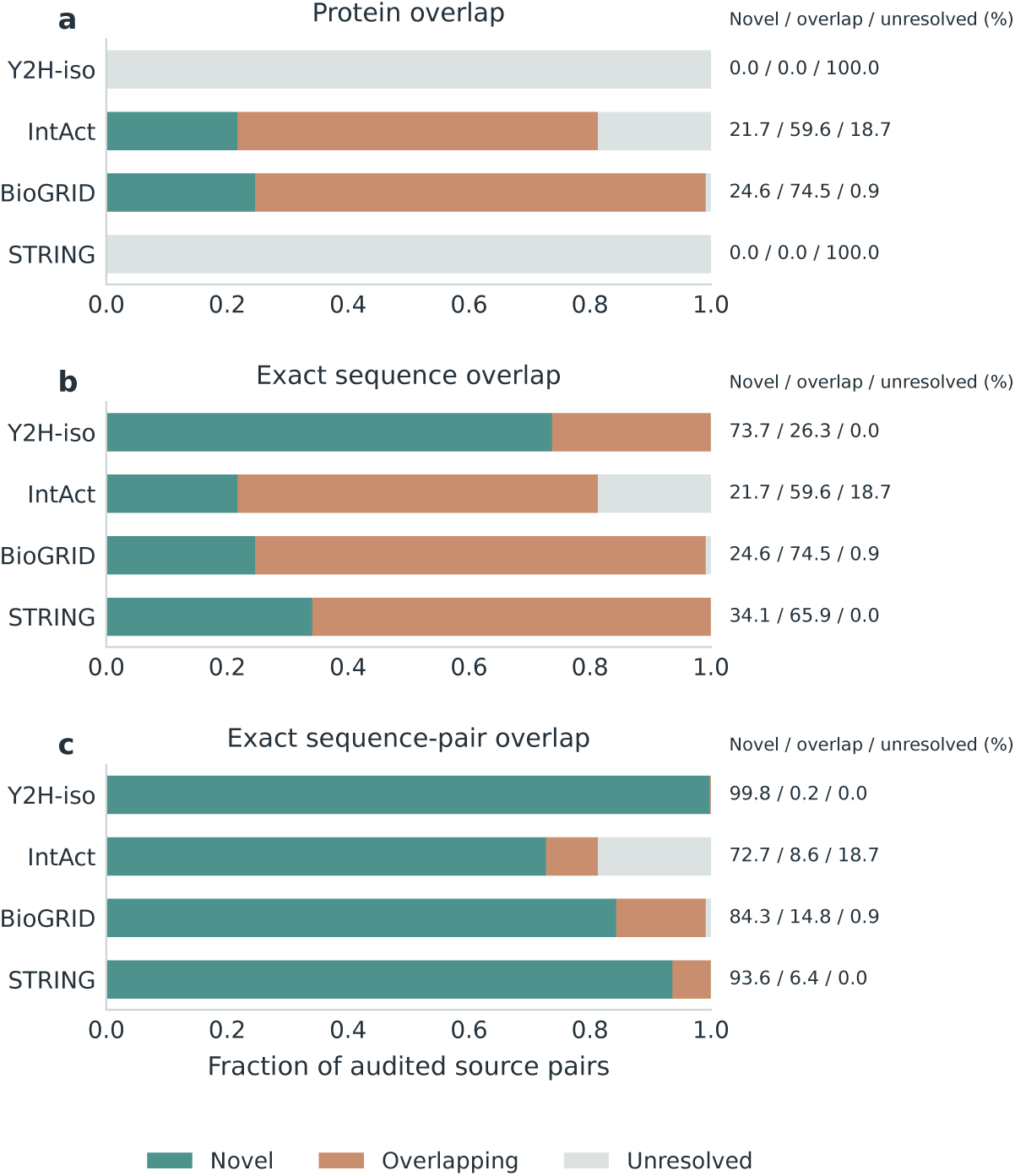
Overlap of external evaluation sources with Gold training and validation data. Protein-identifier, exact-sequence, and exact sequence-pair overlap are separated into novel, overlapping, and unresolved cases. Unresolved denotes unavailable or incomplete identifier mapping and must not be interpreted as biological novelty.

**Figure S11:**
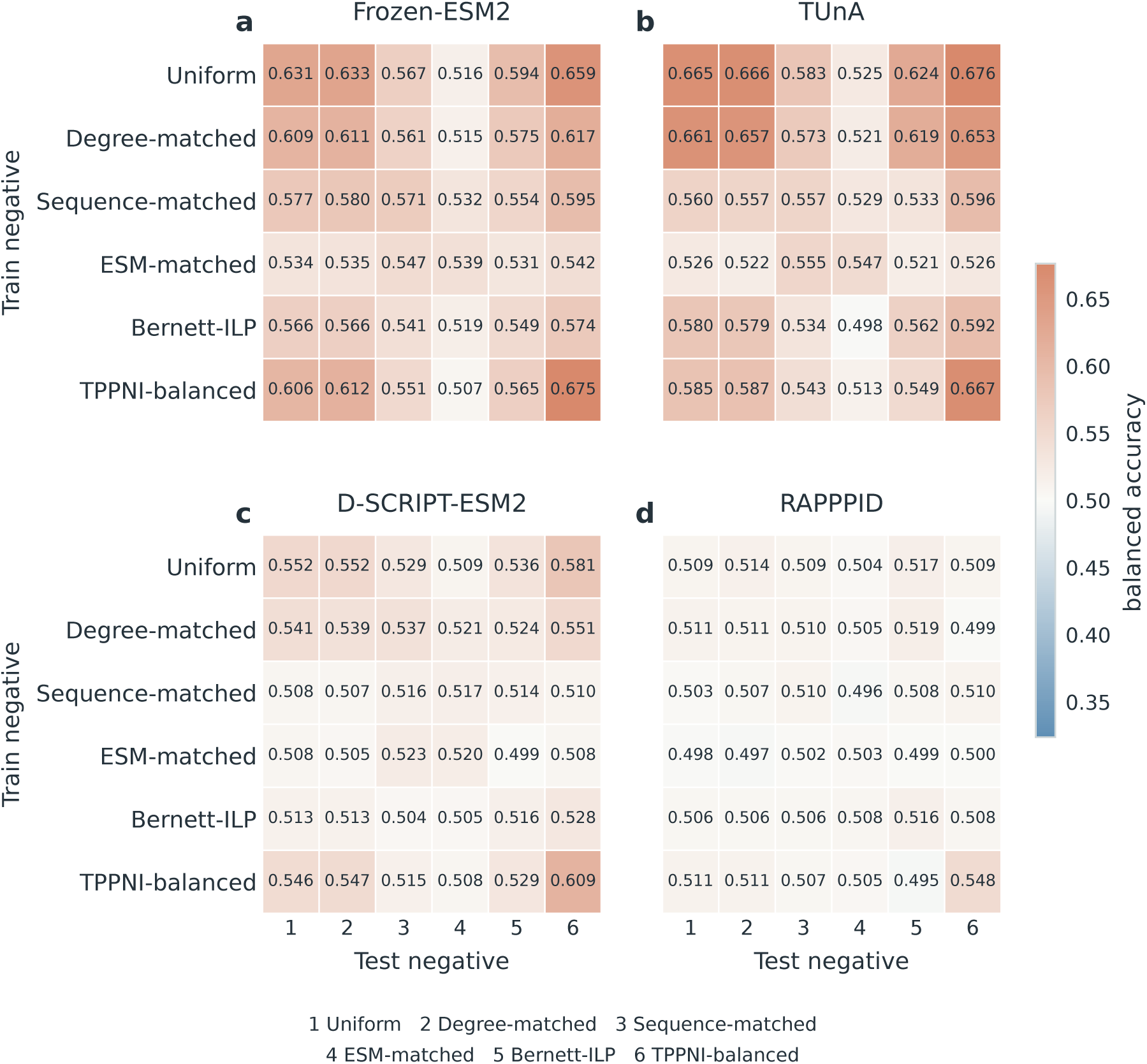
Balanced accuracy across the complete Gold factorial. Thresholds are selected on the matched validation construction and are not re-optimized on test data.

**Figure S12:**
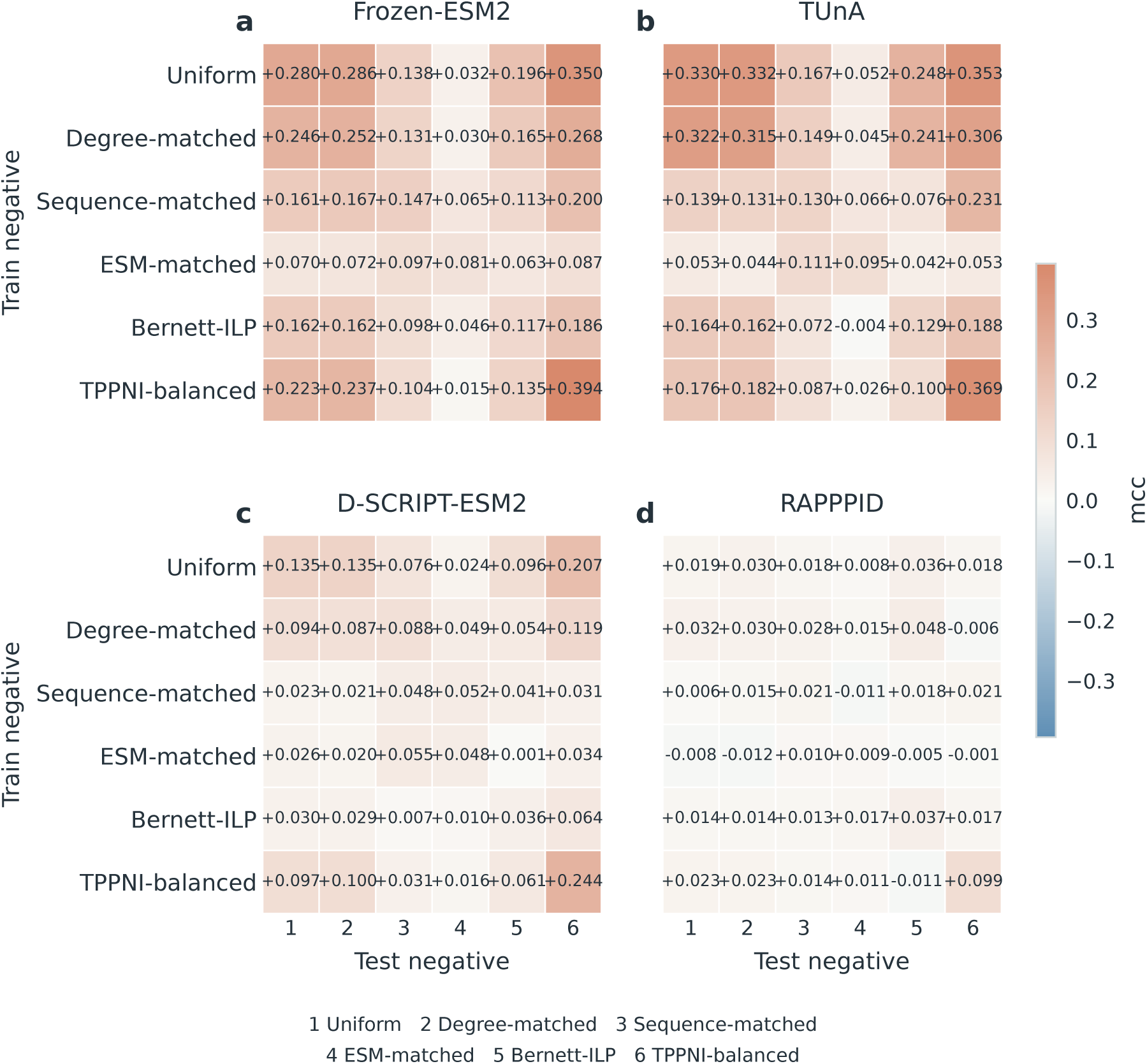
Matthews correlation coefficient across the complete Gold factorial. Thresholds are selected on the matched validation construction and are not re-optimized on test data.

**Figure S13:**
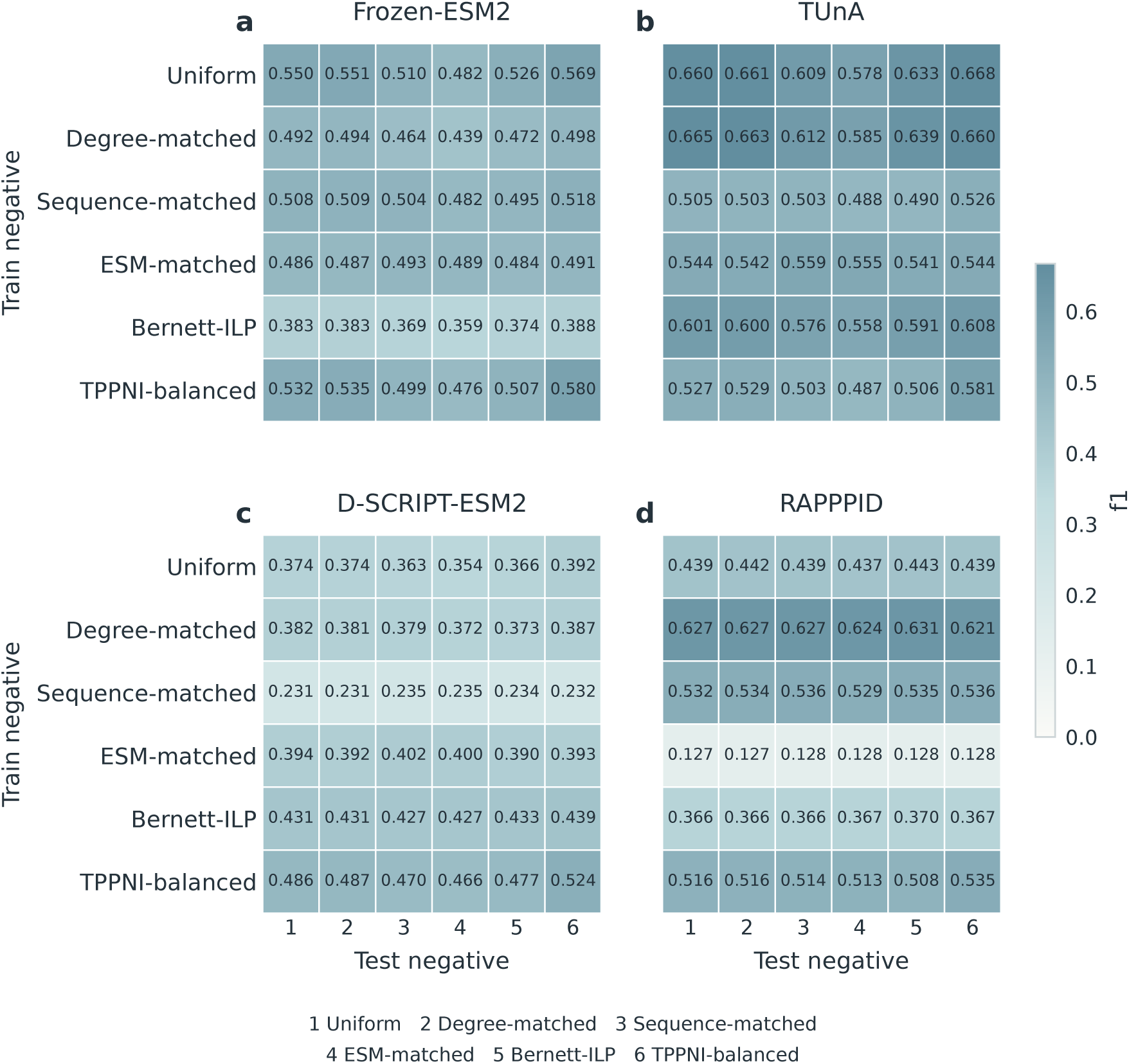
F1 score across the complete Gold factorial. Thresholds are selected on the matched validation construction and are not re-optimized on test data.

**Figure S14:**
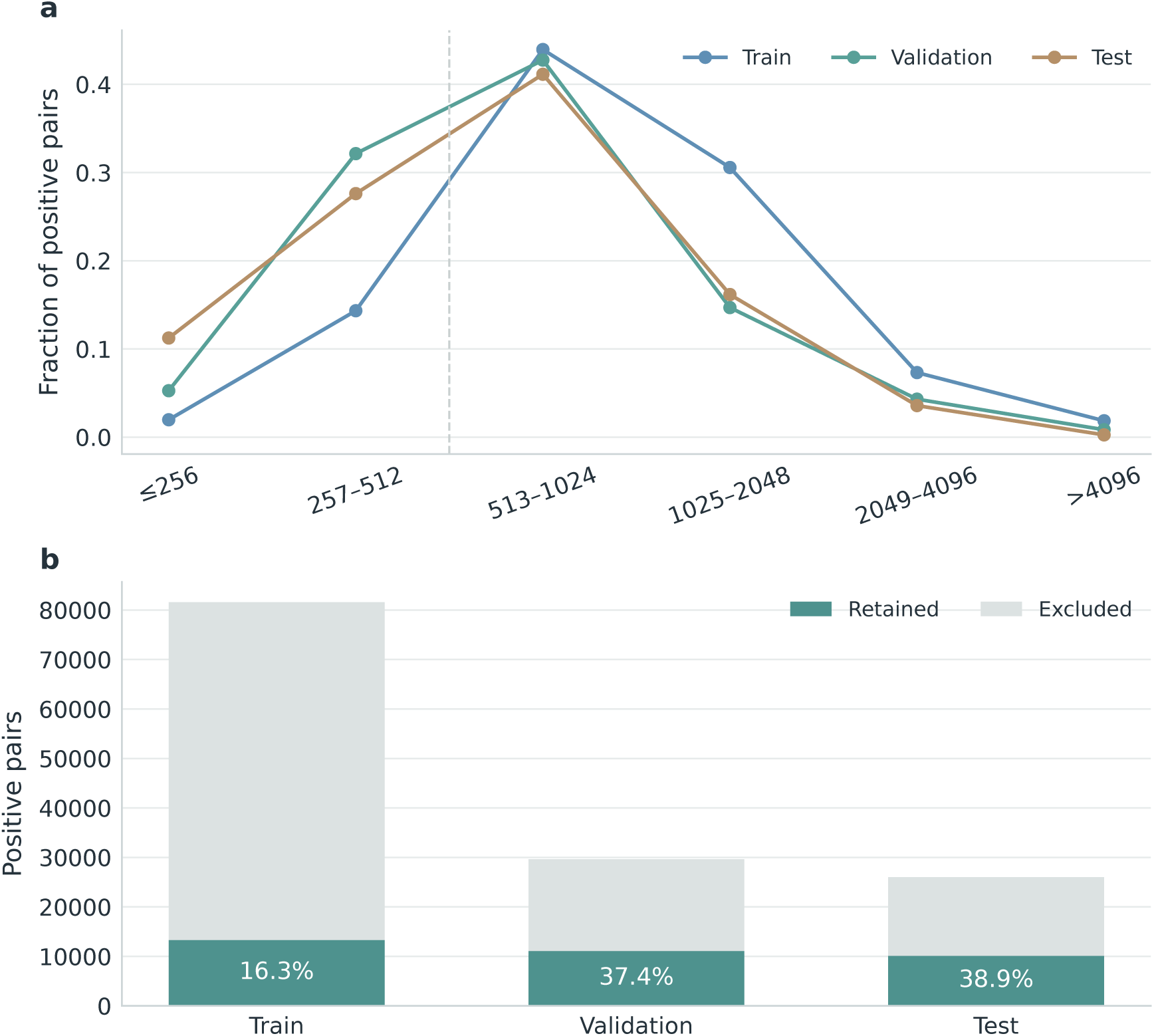
Effect of the common 512-residue compatibility restriction. Panel a shows the distribution of maximum partner length in the original positive pairs for train, validation, and test. Panel b reports retained and excluded positive pairs for each Gold split under the common requirement that both protein sequences contain at most 512 residues.

